# Selective Observation Under Limited Resources Biases Social Inference Through Hysteresis

**DOI:** 10.1101/2025.07.30.667795

**Authors:** Sangkyu Son, Seng Bum Michael Yoo

## Abstract

Despite limited access to others’ actions and outcomes, humans excel at inferring hidden intentions. Given only partial access, how do they decide what to observe, and how does selective observation shape inference? Here, we examined how choosing what to observe can bias the inference about others’ intentions. Participants played a game where they pursued a fleeing target while a computerized opponent acted competitively or cooperatively. Participants overestimated the opponent’s competitiveness after the opponent acted more competitively than expected, whereas no such bias occurred when the opponent was more cooperative than expected. This asymmetry depended on the sequence of events, resembling hysteresis, a form of path dependence observed in physical systems. We found that these biases became stronger when participants chose to observe the opponent instead of their own avatar, and this choice came at the cost of losing precise control over their avatar. Our findings highlight the trade-off in selecting what to observe, as the resulting inference biases propagate differently depending on the interaction history.

## INTRODUCTION

In a social context, one must infer the latent attributes of agents, such as others’ intentions or goals, from a series of observations. For example, when we see an adult following a child, we can infer the adult’s intention, either to prevent the child from falling as a parent or to bypass them as a stranger, by observing a set of actions over time. This inverse process for inferring latent social attributes or others’ mental states through observing others’ actions and following outcomes is called *social inference* (Baker, Saxe, et al., 2009; Cushman, 2024; Jara-Ettinger, 2019). Despite its complexity, humans can seamlessly infer latent social attributes (e.g., roles in the scene) even from animated shapes, as demonstrated in seminal developmental psychology studies (Hamlin et al., 2007; Heider & Simmel, 1944).

In contrast to predicting actions from one’s known goal (i.e., forward problem), social inference is often conceptualized as an inverse problem of deducing one’s latent values or goals from a series of observed actions (Baker, Saxe, et al., 2009; Jara-Ettinger, 2019). From a Bayesian perspective, this can be formalized as inferring the posterior probability of others’ goals given the observed actions (p(goal|action)) (Baker, Saxe, et al., 2009). Similarly, this can be viewed as inverse reinforcement learning, where social inference involves learning others’ state or action value functions through action sequences (Behrens et al., 2009; Jara-Ettinger, 2019). Both emphasize the chain-like inference over time as a Markov Decision Process (MDP), where action sequences serve as the key clues for inferring others’ unobservable goals or values.

While most focus has been on action per se, less attention has been given to the role of observation in sampling those actions. This neglect is attributable to the implicit assumption that one observation could capture all available information, similar to a fully observable Markov decision process (MDP). However, observation is inherently partial and selective—agents do not perceive every action equally, but selectively allocate their gaze to sample socially prioritized aspects of action at each moment. Even among observed information, the socially pertinent attributes can be re-prioritized along the visual pathway (Abassi & Papeo, 2020; McMahon & Isik, 2023; Pitcher & Ungerleider, 2021), including simple shape-based actions (Isik et al., 2017). This suggests that social inference models should expand upon the previous inverse MDP framework and encompass how limited access to action can shape social inference.

Therefore, we re-conceptualize social inference as a bidirectional interaction between the self and others, involving the dynamic unfolding of actions and observations (Figure 1). The key idea is that much of the world beyond one’s own actions and observations is at least partially observable or fully hidden. So, one must strategically allocate the limited observational resources across the environment, others’ actions, and what they observe. For example, in observational learning, the learner attends more to others’ actions to identify intentions relevant to their own problem-solving (Burke et al., 2010; Grabenhorst et al., 2019; Hill et al., 2016) (Figure 1, red arrow), especially when these actions are more socially valuable (Olsson et al., 2020). On the other hand, if observing others provides no particular benefits in learning, one may treat them as part of the uninformative surrounding environment (Gweon, 2021; Jara-Ettinger et al., 2016) (Figure 1, blue arrow). Another example is social gaze–observing what others see (Figure 1, green arrow); more valuable information can be gained by observing how others selectively observe (Dal Monte et al., 2022; Franch et al., 2024; Gao et al., 2010; Luo & Johnson, 2009). Therefore, the proposed framework is expected to integrate residual evidence from previous studies into a more coherent explanation by incorporating shifts in observational focus.

**Figure 1.**
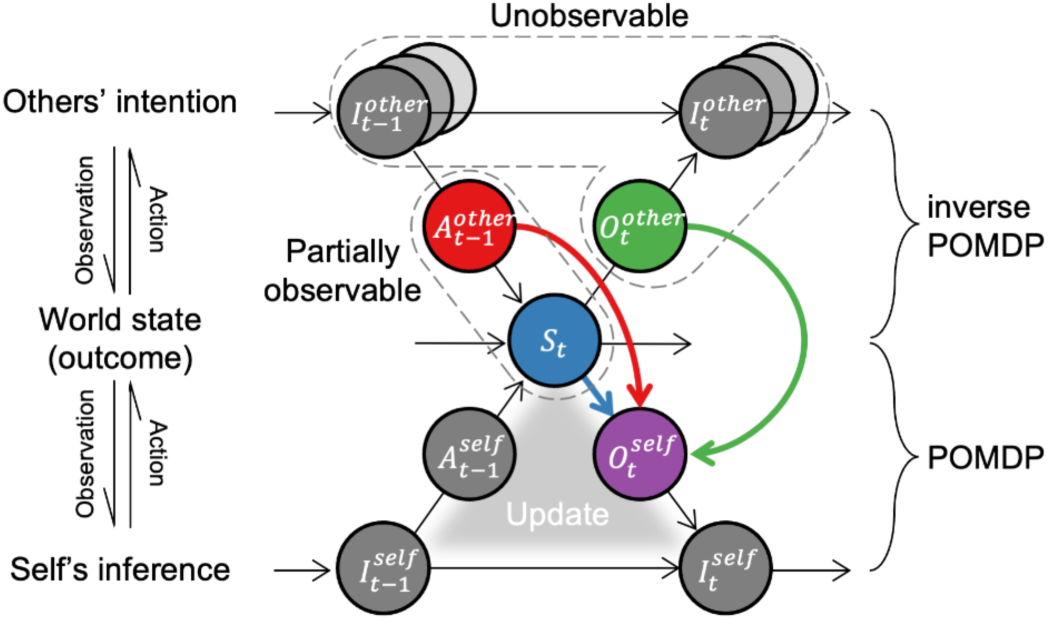
A graphical framework for understanding social inference. Social inference is a bidirectional process between self and others in the world. Both one’s and other’s actions shape the external world as an outcome. Self-world interaction follows a partially observable Markov decision process (POMDP), while other-world interaction follows an inverse POMDP. That is, the agent does not fully know the world state and others’ intentions. When another agent strongly influences the world, the agent prioritizes observing the other’s actions (red) and observations (green) over the rest of the world state (blue).

Within our framework, what would be the consequence of such selective observation in a social context? We reasoned that people update their inferences only when they gain new information from observation; otherwise, their inferences remain unchanged. This suggests that selective observation shapes inference by prioritizing certain social cues over others under a resource constraint. To test this, we designed an interactive task in which participants inferred the hidden intentions of a computer-controlled agent embedded in a dynamic social setting. The agent’s behavior could unexpectedly shift from cooperative to competitive, or vice versa, eliciting positive or negative prediction errors. Because participants could not be monitored simultaneously, they had to allocate their gaze to resolve uncertainty strategically. This design allowed us to investigate how inference biases emerge from the interplay between past social experience and observational focus.

We show that participants’ inferences about the opponent’s intentions were shaped by the recent interaction history—particularly following betrayal. This history dependence was asymmetric: accumulated betrayal led participants to infer greater competitiveness than warranted, whereas comparable experiences of unexpected help failed to induce a similarly strong bias toward cooperative intent. We formalized this asymmetry using an energy landscape framework, revealing that repeated betrayal shifted the attractor basin of the inference dynamics toward competitive interpretations. Eye-tracking data further demonstrated that this shift was followed by a selective allocation of gaze toward the opponent at the expense of monitoring one’s own avatar, reflecting a trade-off between information gathering and control. To probe the mechanism further, we trained a task-optimized recurrent neural network (RNN), which reproduced the asymmetrical inference pattern and revealed that redirecting observational focus could causally reverse its sensitivity to betrayal. These results suggest that selective observation under resource constraints amplifies path-dependent social inference, offering a unified framework that links attention, decision dynamics, and internal models of others.

## RESULTS

### Interactive Social Task To Infer the Hidden Intention of the Opponent

Fifty-four participants (31 males, 21 females; age: 22.7 ± 2.3) played a two-dimensional interactive prey-pursuit game (Figure 2A). Using a joystick, they were required to catch prey that fled from both their avatar and a computerized opponent (Figure 2B). The reward amount decreased as the capture time increased. The opponent’s moment-by-moment movement was determined by a hidden intention parameter (friendliness, F) that balances two conflicting goals: reducing its distance to the prey or the participant’s distance to the prey (Figure 2C). A higher F led the opponent to reduce the distance between the participant and the prey, herding the prey closer to the player. In contrast, a lower F led the opponent to reduce the distance to itself, intercepting the prey before the player. Each trial started with participants watching, on average, a 2.3-second video of three characters interacting, and their eyes were allowed to move freely (inference phase). They were instructed to evaluate the opponent’s hidden intention during the inference phase and then decide whether to increase (i.e., boost) or decrease (i.e., hinder) the opponent’s speed in the subsequent pursuit phase. Strategic decisions were essential since the prey was set to be faster than the participant; boosting a competitive opponent increased the likelihood of failing to capture the prey within the time limit, whereas boosting a cooperative one increased the likelihood of a successful capture.

**Figure 2.**
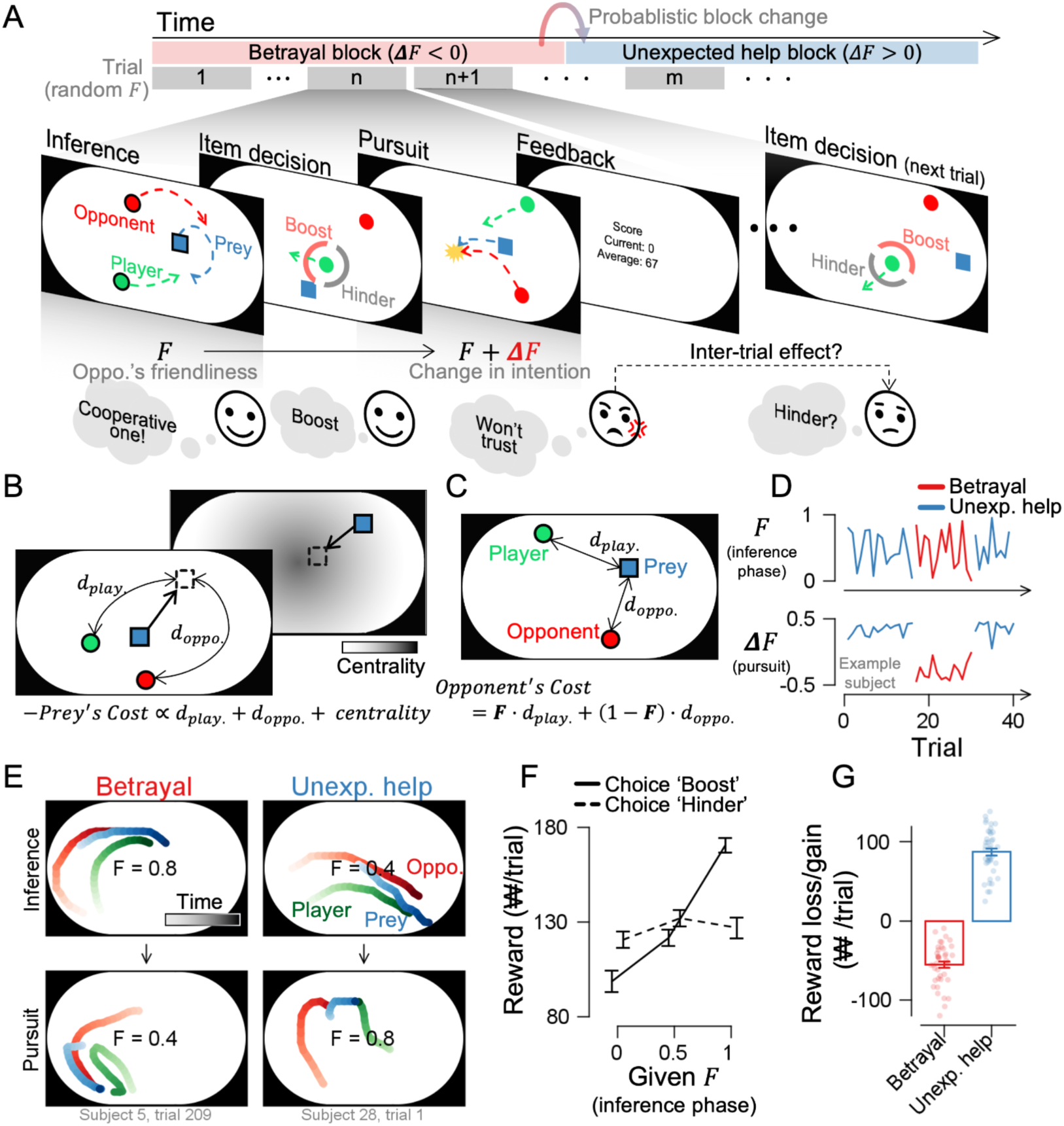
Task schematics of Experiment 1. **(A)** Participants watched videos of their avatar (green) and a computer opponent (red) pursuing prey (blue). The opponent’s hidden intention (F) ranged from competitive (F = 0) to cooperative (F = 1). Participants then decided whether to boost or hinder the opponent’s speed before pursuing prey themselves with a joystick. Unbeknownst to the participants, the F value decreased during the pursuit phase in the betrayal condition and increased in the unexpected help condition. Participants experienced each condition in blocks, with random changes occurring about every 20 trials. **(B)** Prey algorithm: The prey fled from both agents to maximize distance, avoiding walls via a centrality cost. **(C)** Opponent algorithm: At F=1, opponents minimized the distance between the player and the prey (cooperative); at F=0, they minimized the distance between themselves and the prey (competitive). **(D)** During the inference phase, the F value changed randomly, and the direction of ΔF— negative or positive—varied across blocks. **(E)** Example trials. Left: In the betrayal condition, the opponent herded the prey during the inference phase and intercepted it during the pursuit. Right: In the unexpected help condition, the opponent intercepted the prey during the inference phase and herded it during the pursuit. See also Supplementary Videos 1 and 2. **(F)** Boosting a high-F opponent earned higher rewards and hindering a low-F opponent was more beneficial. **(G)** Reward losses from boosting a betraying opponent and gains from boosting an unexpectedly helpful opponent. Error bars represent ±1 SEM, and dots indicate individual participants. ₩ denotes the Korean currency, won.

Unbeknownst to participants, the opponent’s F value changed after their decision (Figure 2D). F value decreased during the pursuit phase compared to the inference phase in the betrayal scenario, whereas it increased in the unexpected-help scenario. In Experiment 1, these scenarios alternated probabilistically. In betrayal, the opponent appeared to herd the prey toward the player in the inference phase but intercepted it in the subsequent pursuit phase (Figure 2E, left panel; Supplementary Video 1). On the other hand, in unexpected help, the opponent chased the prey in the inference phase but herded it toward the player in the pursuit phase (Figure 2E, right panel; Supplementary Video 2). Participants acquired more monetary reward by boosting the opponent with high-F in the inference phase and hindering the opponent with low-F (Figure 2F; choice boost VS. hinder in low-F, mid-F, and high-F, *t*_39_ = −3.168, −2.327, 8.545 with *p* = 0.003, 0.025, 2.035e^-10^, respectively). They earned lower rewards when mistakenly boosting a betraying opponent but gained more when boosting an unexpectedly helpful one (Figure 2G; betrayal and unexpected help, *t*_39_ = −13.407, 18.993, with *p* = 3.331e^-16^, 2.604e^-21^, respectively). These results demonstrate that the interactive social task prompted participants to distinguish between betrayal and unexpected help and adjust their choices accordingly to maximize rewards.

### Asymmetrical history dependence in inference

Humans learn from others’ past actions and intentions, shaping their subsequent social inference (Baker, Goodman, et al., 2009; Baker, Saxe, et al., 2009). Hence, we hypothesized that participants would infer the hidden intentions of the opponent differently, depending on their past interactions, even when observing the behavior with the same F value. Indeed, after experiencing betrayal, participants were less likely to boost the opponent relative to the opponent’s actual F value (Figure 3A and Supplementary Figure 1A; red lines below the unity line; *t*_39_ = −4.665, *p* = 3.576e^-5^). In contrast, after experiencing unexpected help, participants did not show a bias toward boosting the opponent; instead, their choices reflected more accurate predictions of the F values (blue lines overlap with the unity line; *t*_39_ = −0.115, *p* = 0.908). This shows asymmetrical sensitivity, where betrayal caused participants to interpret the opponent’s hidden intentions as more competitive, whereas equivalent unexpected help did not significantly alter their inferences. This asymmetry was independent of sex (Supplementary Figure 1B; for male, *t*_22,betrayal_ = −3.205, *p* = 0.004, *t*_22,unexp. help_ = −1.367, *p* = 0.1854; for female, *t*_16,betrayal_ = - 3.327, *p* = 0.004, *t*_16,unexp. help_ = 1.117, *p* = 0.280) and age (Supplementary Figure 1C; ρ_38,betrayal_ = 0.028, *p* = 0.864, ρ_38,unexp. help_ = −0.135, *p* = 0.408). The asymmetry increased as betrayal experiences persisted: repeated betrayal progressively reinforced the bias toward more competitive inferences (Supplementary Figure 1D; linear trend analysis, *F*_(17, 39), betrayal_ = 18.264, p = 2.205e^-5^; *F*_(17, 39), unexp. help_ = 3.359, p = 0.07). However, the biases after the betrayal experience immediately returned to a neutral state when the betrayal ceased (i.e., at block change) (Supplementary Figure 1E). This indicates that the asymmetrical inference is highly sensitive to the history of changes in betrayal, demonstrating a clear history-dependent characteristic.

**Figure 3.**
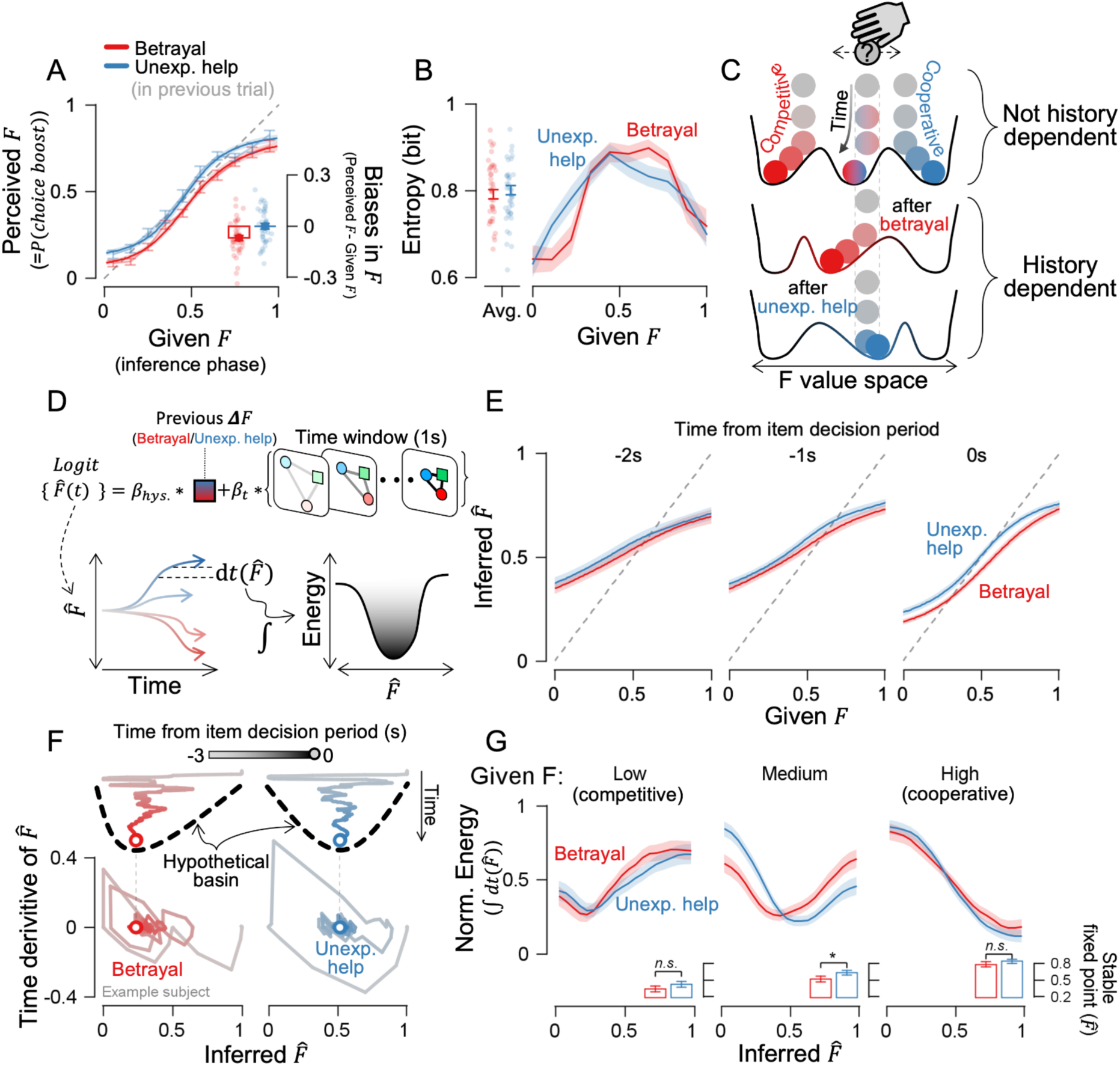
Asymmetrical history dependence in inference. **(A)** The psychometric curve shows the probability of choosing the boost item dropped after betrayal. Bold lines show cumulative Gaussian fits, and faint lines show binned data. The subfigure shows biases in their decisions, calculated as perceived F minus given F. **(B)** Shannon entropy of given F showing uncertainty in participant’s choice. **(C)** F-value inference during the inference phase resembles a ball rolling on a landscape with basins. If prior history alters the landscape, identical stimuli would settle at different points. **(D)** Schematic of inference landscape reconstruction: moving-windowed trajectories and the previous trial’s ΔF were used in an autoregressive logistic regression model. Temporal changes in model-estimated 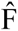 were accumulated to build the energy landscape. **(E)** The psychometric curve, derived from the logistic regression model, shows how inference evolves over time until the decision phase begins. **(F)** Example subject’s phase portrait of inferred 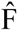 value after betrayal and unexpected help. A positive 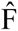 derivative on the y-axis indicates an increase in 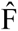 at the next time point on the x-axis, and vice versa. **(G)** Reconstructed energy landscapes, split into trials with low (F < 0.33), medium (0.33 < F < 0.67), and high F value (F > 0.76). The lowest energy points (stable fixed points) show where inferences would stabilize in the subfigures. Shaded ribbons and error bars represent ±1 SEM, and stars indicate statistical significance (*, p < 0.05; n.s., not significant).

Given the asymmetrical history dependence in social inference, we examined which aspect of past interaction contributes to this effect. One possibility is that betrayal caused greater monetary loss than the gain from unexpected help. However, the opposite was true: the gain was larger in the unexpected help condition (Figure 2G; betrayal VS. unexpected help, *t*_39_ = 6.211, p = 2.634e-7), indicating that reward value differences do not drive the asymmetry. Another possibility is that each type of experience resulted in a different level of uncertainty in the upcoming trial. We converted choice probabilities into Shannon entropy (see Materials and methods for details), a measure used for quantifying the internal uncertainty of participants choosing certain options (Kepecs et al., 2008; Meyniel et al., 2015; Wang et al., 2023). On average, Shannon entropy was similar between betrayal and unexpected help trials (Figure 3B, left panel; *t*_39_ = 0.659 with *p* = 0.513). However, when entropy was analyzed at each level of the given F value separately, participants’ confidence in their choices varied according to their interaction history (Figure 3B, right panel); the entropy was higher following betrayal than unexpected help when F < 0.5, but this pattern reversed when F > 0.5, with higher entropy following unexpected help (Cluter-based permutation t-test, betrayal < unexpected help, *t*_39_ = 2.039, 2.801, with *p* = 0.003 for both, at F = 0.15 and 0.25, respectively; betrayal > unexpected help, *t*_39_ = - 2.716 with *p* = 0.048 at F = 0.65). Additionally, decision times were similar across conditions (Supplementary Figure 1F; betrayal vs. unexpected help, all *p* > 0.05), indicating that differences in entropy were not due to longer decision durations but rather reflected participants’ internal uncertainty about their choices. These suggest that prior experiences altered the overall internal uncertainty of participants’ inference and shifted where participants felt most certain.

### The energy landscape captures the asymmetry

Uncertainty in participants’ choices relates to history-dependent inference, but how this uncertainty translates into inferential bias remains unclear. We approached this question using the framework often referred to as an energy landscape and interpreted uncertainty reduction as the system settling into an energetically stable state (i.e., stable fixed point) (Wang et al., 2023) (Figure 3C). The energy landscape framework transforms uncertainty into an intuitive geometric representation and offers predictive power, as decisions follow subject-specific dynamics shaped by uniquely formed landscapes.

We reconstructed the energy landscape by tracking how inference stabilized over time using an autoregressive logistic regression model (Figure 3D). Since the opponent’s appearance provided no clues about their hidden intention, participants had to rely solely on the temporal patterns of the three characters’ movement and prior experiences of betrayal or unexpected help. Thus, the model incorporated two key parameters: β_t_, which captures how well participants infer the opponent’s intentions from the trajectories of the three characters during the inference phase, and β_hys_, which regulates how past experiences of betrayal or unexpected help influence inference bias. In sum, β_t_ primarily alters the steepness of the psychometric curve, while β_hys_ primarily modulates the magnitude of inference bias (Supplementary Figure 2). We then mapped the inference landscape based on how inference stabilizes across levels of perceived F-value space (Wang et al., 2023) (see Materials and methods for details).

We first examined when history-dependent inference arises—whether it stems from initial conditions before inference begins, or from dynamic changes as inference unfolds. The model supported the latter hypothesis. Asymmetric biases gradually emerged in the psychometric curves, with betrayal leading to a lower F value compared to unexpected help (Figure 3E), mirroring the final decision pattern shown in Figure 3A. Moment-by-moment changes in inference—known as a phase portrait—visualized this more clearly; while early fluctuations appeared similar, inference eventually settled down to a more competitive interpretation after betrayal (Figure 3F). This suggests that asymmetrical history dependency arose from changes in inference dynamics rather than from an initial baseline condition shift.

Which aspect of the inference dynamics changed after betrayal? We considered two possibilities. First, the convergence point becomes more deeply embedded after betrayal than before, reflecting a higher degree of confidence in the decision. This resembles a steeper or deeper valley—harder to escape and more stable—which is known to reflect stronger decision confidence (Atiya et al., 2019, 2021; Wang et al., 2023). Second, the convergence point itself may shift laterally, meaning the bottom of the valley moves. The reconstructed energy landscape supported the convergence shift, especially when the opponents behaved ambiguously. The points of lowest energy shifted significantly toward lower inferred F values after betrayal, especially when participants encountered an opponent displaying ambiguous behavior with mid-range F values (Figure 3G; betrayal vs. unexpected help across low-F, mid-F, and high-F; *t*_39_ = 1.605, 2.261, 0.977; *p* = 0.116, 0.029, 0.334, respectively). In contrast, the steepness or depth of attractor basins remained consistent across different prior experiences (Supplementary Figure 3; across F-value ranges, *F*_(2,78)_ = 7.416, 0.792, with *p* = 8.375e^-4^, 0.455; across betrayal VS. unexpected help, *F*_(1,78)_ = 0.002, 0.046, with *p* = 0.966, 0.831; and interaction, *F*_(2,78)_ = 0.354, 2.036, with *p* = 0.702, 0.134, depth and steepness, respectively). The result indicates that history-dependent inference arises not from changes in the degree of confidence (i.e., deeper basin), but rather from the shift in decision criteria (i.e., shift of basin).

### Observational focus shifts adjust inference

We wondered how previous experiences shift where the final inference ultimately converges. We hypothesized that how people gather information after betrayal could systematically shift where the final inference converges. When we analyzed eye traces to test this hypothesis, participants more often fixated on the opponent during betrayal, while they focused on their own avatar during unexpected help (Figure 4A). By comparing the distance between the gaze center and each agent on the screen, we found that during the initial 1.5 seconds of pursuit, participants focused on their own avatars to locate themselves in both betrayal and unexpected help contexts. However, even when F values were matched, participants shifted their gaze to the opponent shortly after the initial 1.5 seconds in betrayal trials, whereas their gaze remained focused on their own avatar during unexpected help trials (Figure 4B; cluster-based permutation test, betrayal VS. unexpected help, overall trials, *p* < 0.001 after 1.08 seconds from the pursuit phase onset; Supplementary Figure 4A, F values matched, *p* < 0.001 after 1.17 seconds from the pursuit phase onset). Ultimately, how observational resources are allocated to the opponent predicted inference in subsequent trials. When participants observed the opponent more closely, they were more likely to infer them as a competitor in later trials (Figure 4C, *p* = 0.007 from 0.98 to 1.70 and *p* = 0.023 from 1.90 to 2.32 seconds after the onset of the pursuit phase). Similarly, longer observation durations also increased the likelihood of inferring competitive intent (Figure 4D, t_39_ = −2.478 with *p* = 0.02). The proportion of saccadic eye movements did not differ between contexts, indicating that rapid shifts in eye position from one agent to another were not the primary driver of the observational shift (Supplementary Figure 4B; *p* > 0.05 for all time points). This suggests that the opponent’s betrayal behavior led participants to shift their gaze to the opponent to gather additional information, which in turn influenced their subsequent inference process later.

**Figure 4.**
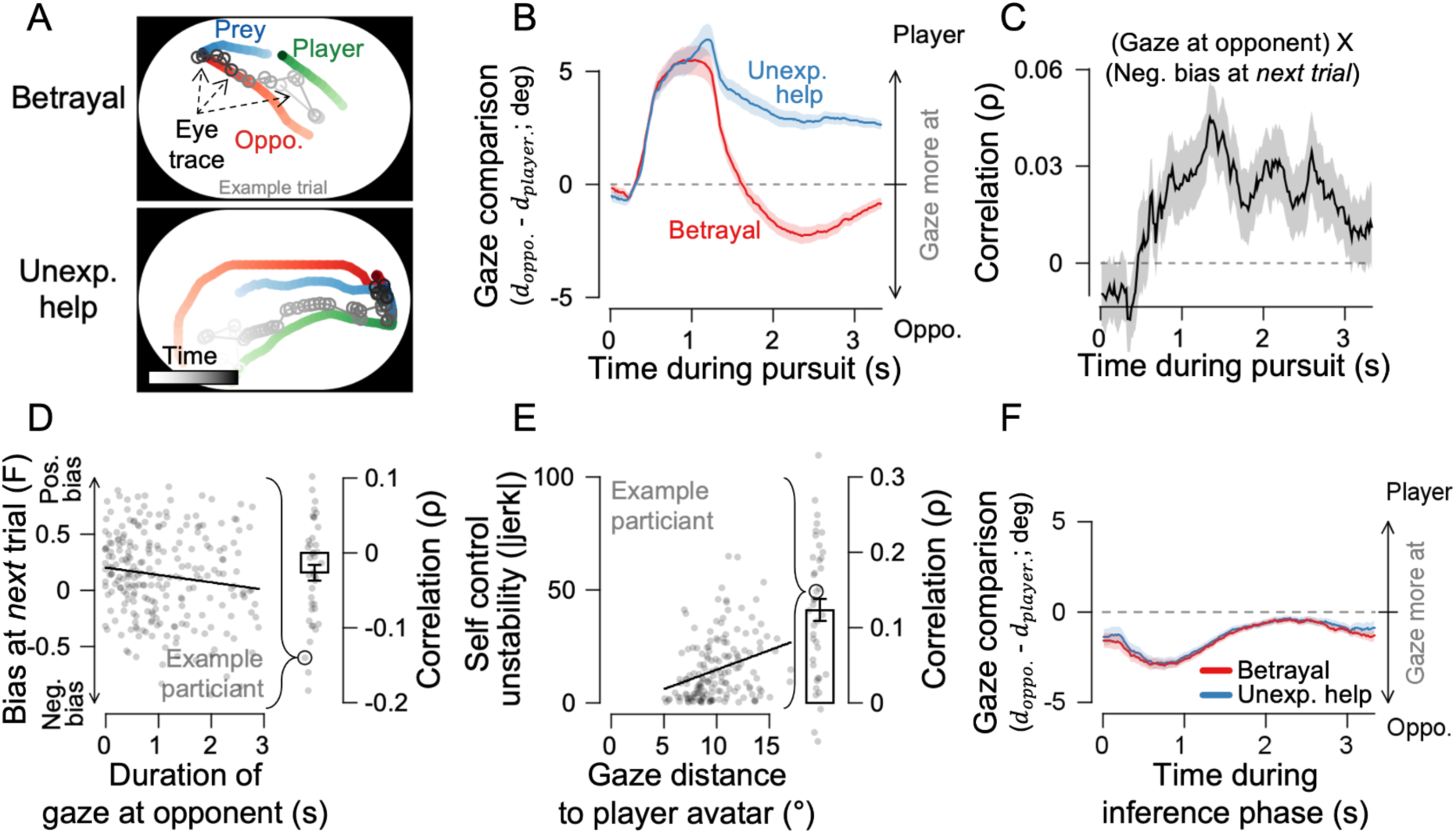
Observational focus shifts adjust inference. **(A)** Eye-tracking examples show gaze shifts toward the opponent during betrayal trials but not during unexpected-help trials in the pursuit phase. **(B)** Comparison of the distance from the gaze center to the opponent’s avatar (d_oppo._) and the player’s avatar (d_play._). **(C)** Spearman’s correlation between gaze at the opponent and competitor biases in perceived F in the subsequent trial. **(D)** Longer gaze durations toward the opponent correlate with biases in perceived F in the subsequent trial. **(E)** A farther gaze from oneself correlates with lower stability (higher jerk) in the self’s avatar movement. **(F)** Gaze comparison during the inference phase period. Error bars and shaded ribbons indicate ±1 SEM, with dots representing individual trials.

We wondered what participants might sacrifice to obtain additional information. We reasoned that acquiring additional information would come at the expense of precise control of the participant’s own avatar due to reduced time spent fixating on it. Indeed, shifting gaze away from one’s avatar led to decreased control stability, as indicated by an increased jerk, reflecting a more abrupt acceleration in self-avatar trajectories (Figure 4E; *t*_39_ = 19.950, *p* = 4.562e^-22^). However, when the necessity for control is removed during the observation phase in the following trial, the gaze pattern did not differ between the two past histories and mostly gazed at the opponent where the most information is (Figure 4F; *p* > 0.05 for all time points). Overall, these trade-offs suggest that participants prioritized what to observe based on cost; the opponent’s betrayal was more costly than losing some control over their own avatar. This cost-driven observation became a natural source of information asymmetry. That is, by shifting their gaze to the opponent, participants gained more information about unexpected competitiveness; otherwise, they learned nothing new. Our results suggest that this difference in information from selective observation eventually biased the later inferences.

### Observation target change counteracts inferential bias

To uncover the underlying causal role of observation in the inference process, we developed task-optimized RNNs and causally manipulated the network (Figure 5A). First, the RNN model was trained to predict a scalar value of a given F value from the movies during the inference phase without using participant choices (i.e., task-optimized RNN). This step was to obtain the neutral inference machine. In the second step, we examined how the artificial network adapted to the movies during the pursuit phase, where the participants experienced either betrayal or unexpected help scenarios. Betrayal and unexpected-help scenarios led to different types of discrepancy when fed into the neutral inference machine from step 1: the inferred 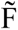 would deviate negatively from the expected F (given from the inference phase) in betrayal scenarios, and positively in unexpected-help scenarios. We introduced a free parameter, ꞵ_bet_, to balance between betrayal and unexpected help, assessing the network’s sensitivity to betrayal. The network also adjusted which inputs from the three characters to prioritize in the next training epoch, assuming that changes in the RNN’s input weights could serve as a proxy for humans’ selective observations. Using the neutral inference machine from the first step as a baseline, we repeated the second step to explore how inferences diverged based on the RNN’s distinct adaptation.

**Figure 5.**
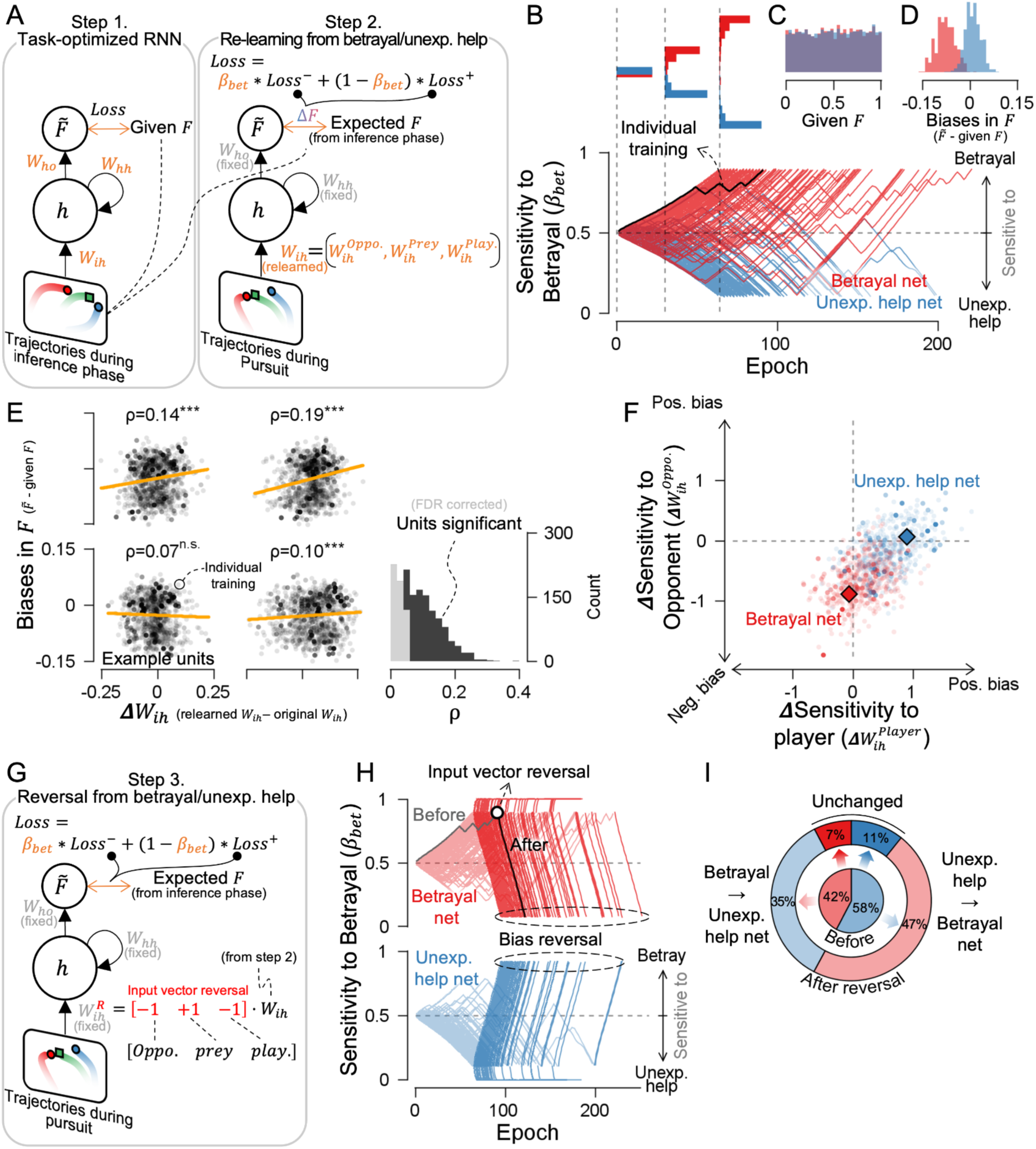
A task-optimized recurrent neural network (RNN) demonstrated that observation plays a causal role in shaping inferential bias. **(A)** RNN training. Step 1: Train the model to predict the F value from the trajectory movie during the inference phase. Step 2: Re-train the model using the movie during the pursuit phase under betrayal or unexpected help conditions. Only the input weights W_ih_ for the opponent, prey, and player were updated while keeping other parameters fixed. β_bet_ balances sensitivity to betrayal and unexpected help. Orange indicates trainable parameters. **(B)** β_bet_ over training epochs in Step 2. Each line represents a separate re-learning from Step 1, categorized as either betrayal-sensitive or unexpected-help-sensitive if β_bet_ exceeded 0.9 or dropped below 0.1. The upper subfigures show diverging distributions over time. **(C)** Distributions of input F value. **(D)** Output biases in inferred 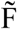 value. **(E)** Changes in input weights for most RNN units during Step 2 correlated with biases in inferred 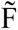 value. Left: Four example units. Right: Correlation distribution across all units. Dots representing individual Step 2 training sessions. **(F)** Input weight changes for opponent and player movement trajectories. Negative changes shift the inferred 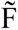 value toward a competitive interpretation, while positive changes shift it toward a cooperative one. **(G)** Step 3: Input weights learned in Step 2 were sign-reversed, and only the betrayal sensitivity parameter β_bet_ was re-trained to test the effect. **(H)** Learning traces of β_bet_ before (Step 2, faint lines) and after sign reversal (Step 3, bold lines). **(I)** Most betrayal-sensitive networks switched to unexpected-help-sensitive networks, and vice versa, after reversal.

Simulations demonstrated that inference networks bifurcated into two categories: those sensitive to betrayal or unexpected help (Figure 5B). The input F values used for training were comparable between the betrayal-sensitive and unexpected-help-sensitive networks (Figure 5C), but the model output from betrayal-sensitive networks showed an inference bias toward negative F values, whereas the unexpected-help-sensitive networks did not (Figure 5D). This asymmetrical pattern of inferential bias mirrors human participants’ responses to betrayal and unexpected-help scenarios, as illustrated in the subfigure of Figure 3A. Changes in input weight correlated with inferential bias; a decrease of the input weight led to a corresponding inference bias towards perceiving the opponent as a competitor (Figure 5E; 65.3% of significant units, false-discovery-rate corrected; average correlation, ρ = 0.121 ± 0.002). Importantly, the betrayal-sensitive network reduced the input weight of opponent movements while keeping the input weight of player movements unchanged, leading to an inference bias toward a negative F value (Figure 5F; permutation test, *p_player_* = 0.532, *p_opponent_* = 0.020). On the other hand, the weight of the unexpected-help-sensitive network remained F value unchanged (*p_player_* = 0.948, *p_opponent_* = 0.435). These RNN results demonstrate that reallocating observational resources to the opponent’s movement mainly drives the inferential bias toward negative F values, revealing the source of the asymmetrical bias observed in human participants.

Given that inference depends on observation, we wondered whether shifting observational focus alone could significantly alter it. We therefore reversed the opponent and player weights—serving as a proxy for observational input—in the task-optimized RNN, and examined changes in betrayal sensitivity (ꞵ_bet_), while keeping all other parameters fixed. (Figure 5G). We found that reversing the signs of input weights, without altering the network structure, was sufficient to reverse sensitivity. Networks previously sensitive to betrayal became sensitive to unexpected help, and vice versa (Figure 5H), indicating that bias reversal occurred frequently across networks (Figure 5I). These reversal results confirm that allocating observational resources causally contributes to inferential bias, but also demonstrate that such bias can be mitigated even when observational resources are re-allocated.

### Hysteresis in social inference

We wondered whether inference would immediately become unbiased once betrayal stopped, or whether it would resemble hysteresis in physical systems—exhibiting lingering effects shaped by the trajectory of past experiences (Figure 6A). We assumed that if the amount of betrayal of the current trial’s one has increased than the previous trial (i.e., more negative ΔF), participants are more likely to interpret the opponent as competitive, compared to when the error has decreased. Hysteresis rejects three alternative possibilities (Figure 6B): (1) inferential bias is independent of recent experiences; (2) all recent experiences have an equal and symmetrical effect on bias; and (3) even if asymmetrical, there is no path dependence—meaning no distinction between increasing and decreasing phases of betrayal. To test for hysteresis in inference, in the subsequent experiment (Experiment 2), we introduced a monotonic increase or decrease in the opponent’s intention change (ΔF) between the inference and pursuit phases, alternating these phases to create a zigzag pattern of ΔF (Figure 6C).

**Figure 6.**
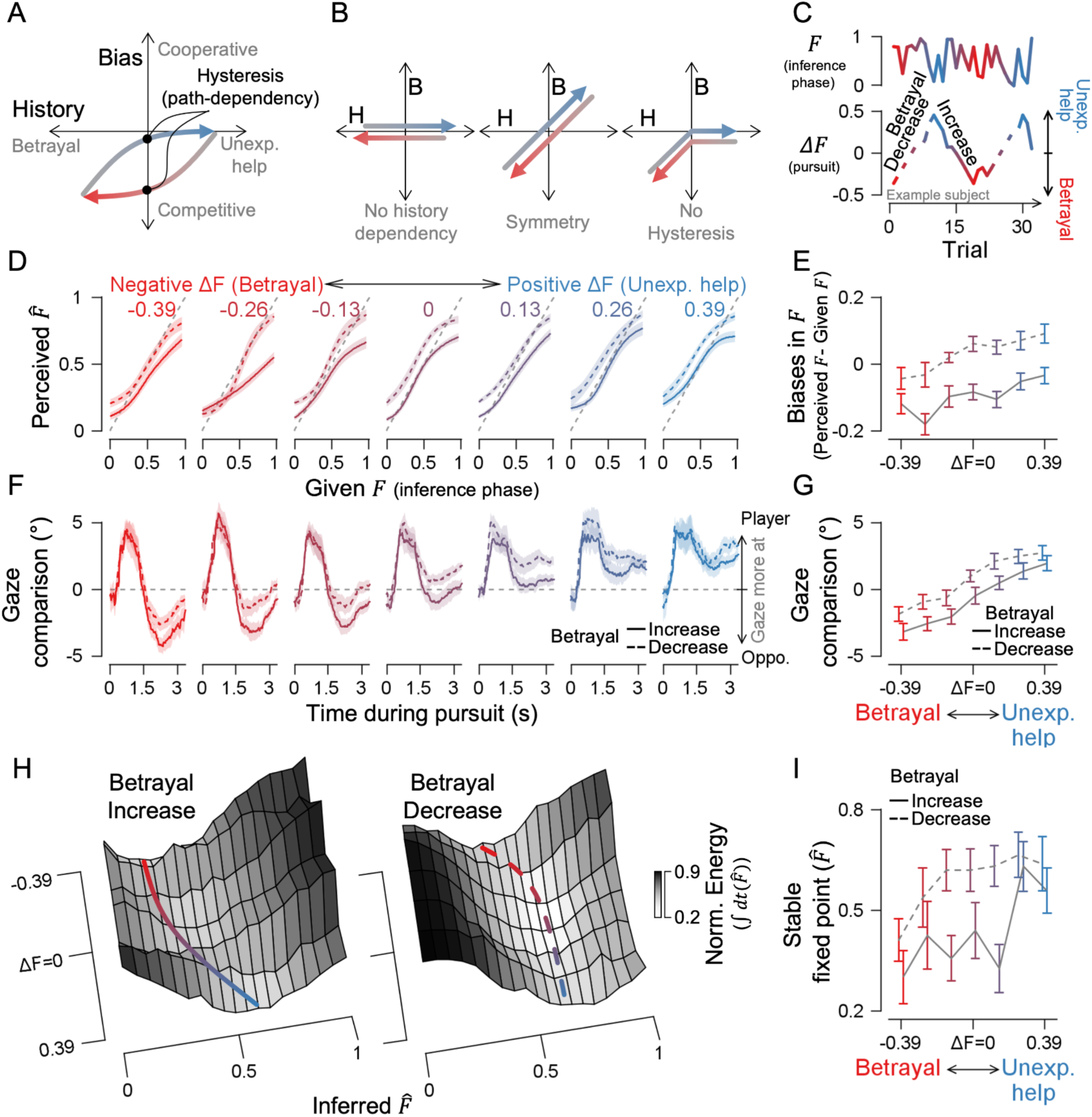
Hysteresis in inference in Experiment 2. **(A)** Null hypothesis: If the order of experience influences inference (i.e., hysteresis exists), a gradual increase and decrease in betrayal strength should bias inference differently. **(B)** Alternative hypotheses. Left: biases are history-independent; Middle: Biases depend on past experiences and each experience holds equal, symmetrical value; Right: biases depend asymmetrically on betrayal but not on the exact order of events (i.e., not path-dependent). **(C)** During the inference phase, the F value randomized while ΔF alternated in a zig-zag pattern, shifting between increasing and decreasing betrayal. **(D-E)** Psychometric curves and inferential biases by levels of betrayal or help. **(F-G)** Gaze comparisons between player and opponent by levels of betrayal or help. **(H)** Energy landscapes for moderate F value (0.33 < F < 0.67); basin lines mark the approximate lowest points. **(I)** Phase diagram showing where inference settles into lower-energy stable points. Shaded ribbons and error bars indicate ±1 SEM. Solid lines show an increasing betrayal phase, and dashed lines decreasing betrayal phase.

As in Experiment 1, stronger betrayal experiences in previous trials led to a greater bias toward perceiving the opponent as a competitor, while unexpected help did not create a significant bias (Figure 6D-E). However, this effect became more pronounced when the degree of betrayal increased across trials, compared to when it decreased (increase VS. decrease; *t_13_* = −2.347, −3.677, −3.739, −5.722, −6.713, −3.879, −5.180, with p =0.035, 0.003, 0.003, 7.019e^-5^, 1.439e^-5^, 0.002, 1.768e^-4^, respectively for the seven bins ranging from the strongest to the weakest betrayal). A similar pattern of gaze behavior was observed, consistent with the findings from Experiment 1 (Figure 6F-G). Participants generally focused on the opponent during betrayal and their own avatar during unexpected help but fixated on the opponent more closely during the phase of betrayal increase than during the phase of betrayal decrease (increase VS. decrease; *t_13_* = −3.042, −4.889, −4.644, −4.604, −7.836, −3.466, −2.122, with p = 0.009, 2.957e^-4^, 4.592e^-4^, 4.934e^-4^, 2.801e^-6^, 0.004, 0.053, respectively for the seven bins ranging from the strongest to the weakest betrayal). This presents an interesting contrast, showing that even with the same experience of an opponent behaving with ambiguous intention, its impact on observation and subsequent inference differs depending on whether the degree of betrayal increased or decreased.

Hysteresis manifested as structural changes in the inference landscape, depending on whether the degree of betrayal was rising or falling. When we reconstructed the landscape from trials where the opponent’s intention was ambiguous (i.e. mid-F range), the stable fixed points with the lowest energy significantly differed between the phases of betrayal increase and decrease, especially when ΔF was nearly zero at the latest trial (Figure 6H-I; betrayal increasing VS. decreasing; *t_13_* = –2.337 and −2.623, with *p* = 0.036 and 0.02 for ΔF = −0.13 and 0.13, respectively, while the rest showed *p* > 0.05). On the other hand, the inference landscape in trials with low or high F-values did not differ between increasing and decreasing phases of betrayal (Supplementary Figure 6; only at ΔF = 0.26 and high-F, *t_13_* = –2.503 with *p* = 0.026, while the rest showed *p* > 0.05). This suggests that stopping the betrayal will not immediately restore neutral inference due to its momentum when the degree of betrayal continues to increase. Instead, our results imply that a sustained accumulation of unexpected help experiences or drastic changes in observational focus might be necessary for restoration.

## DISCUSSION

Like other forms of inference, inferring others’ intentions is subject to bias, particularly as interactions unfold over time in partially observable environments. Here, we demonstrate such biases using an interactive prey-pursuit task involving an opponent with hidden intentions, interpreted through the lens of an energy landscape framework. The observed bias exhibited several notable properties. First, it was history-dependent: past interactions influenced current inferences, with the most potent effects occurring under high uncertainty. Second, the history dependence was asymmetric—participants showed more substantial biases following betrayal than unexpected help. Third, it exhibited hysteresis: the effect of betrayal history was greater when betrayal was an increasing phase than when it was a decreasing phase. We found that how participants allocated their observational resources shaped this bias, while balancing the trade-off between acquiring information about the opponent and maintaining precise control of the self-avatar. Causal intervention using a task-optimized recurrent neural network further confirmed that shifting the focus of observation can reverse inference biases.

What are the functions of observation in social inference? In many situations, uncertainty challenges effective decision-making, especially when insufficient sampling misaligns external conditions with internal models (Berkes et al., 2011). In social contexts, this challenge is amplified: individuals often lack direct access to others’ internal states and must selectively decide what to observe or sample (Figure 1). Traditionally, such uncertainty in social inference is thought to be resolved through high-level cognitive processes—such as adopting others’ perspectives or simulating their experiences (FeldmanHall & Nassar, 2021; FeldmanHall & Shenhav, 2019), or by planning to reduce future ambiguity (Ho et al., 2022). However, our results suggest that shifting gaze serves as a simple yet effective way to reduce uncertainty during social inference. More importantly, we emphasize that observation is selective and thus costly. This raises the question of how to reallocate limited observational resources to reduce specific aspects of social uncertainty. We suggest that a simple shift of gaze determines how the limited observational budget is allocated across social cues, leaving unobserved aspects without resources. For instance, when you concentrate on reading the audience’s polite smiles, you might miss their hand gestures that signal boredom.

While decision-making models from non-social domains have provided valuable starting points for understanding social inference, they may capture only certain aspects of complexity in our result. For instance, a drift-diffusion model (DDM) might account for the current asymmetrical inferential bias by accelerating threshold-crossing after betrayal (Cushman, 2024; Gates et al., 2021; Gold & Shadlen, 2007; Tump et al., 2020) (Figure 7A), while a Bayesian model might explain it by assigning stronger prior beliefs to betrayal than to unexpected help (Baker et al., 2017; Diaconescu et al., 2014; Pisauro et al., 2022; Trudel et al., 2023) (Figure 7B). The canonical form of both models predicts reduced variability after betrayal compared to unexpected help. However, our data show no such difference in decision time or choice variability (Figure 3B and Supplementary Figure 3). To address this discrepancy, we adopted a dynamical systems perspective. A geometric structure in the inference energy landscape intuitively explains why asymmetric bias emerges only when betrayal increases (Figure 7C). This framework also explains why competitor bias is difficult to reverse once it has formed. Reversing it demands either consistent, strongly cooperative behavior from the opponent (green arrow) or redirecting observation toward the self (purple arrow). By embedding the influence of prior states, the dynamical system framework reveals the temporally intricate and path-dependent nature of social cognition.

**Figure 7.**
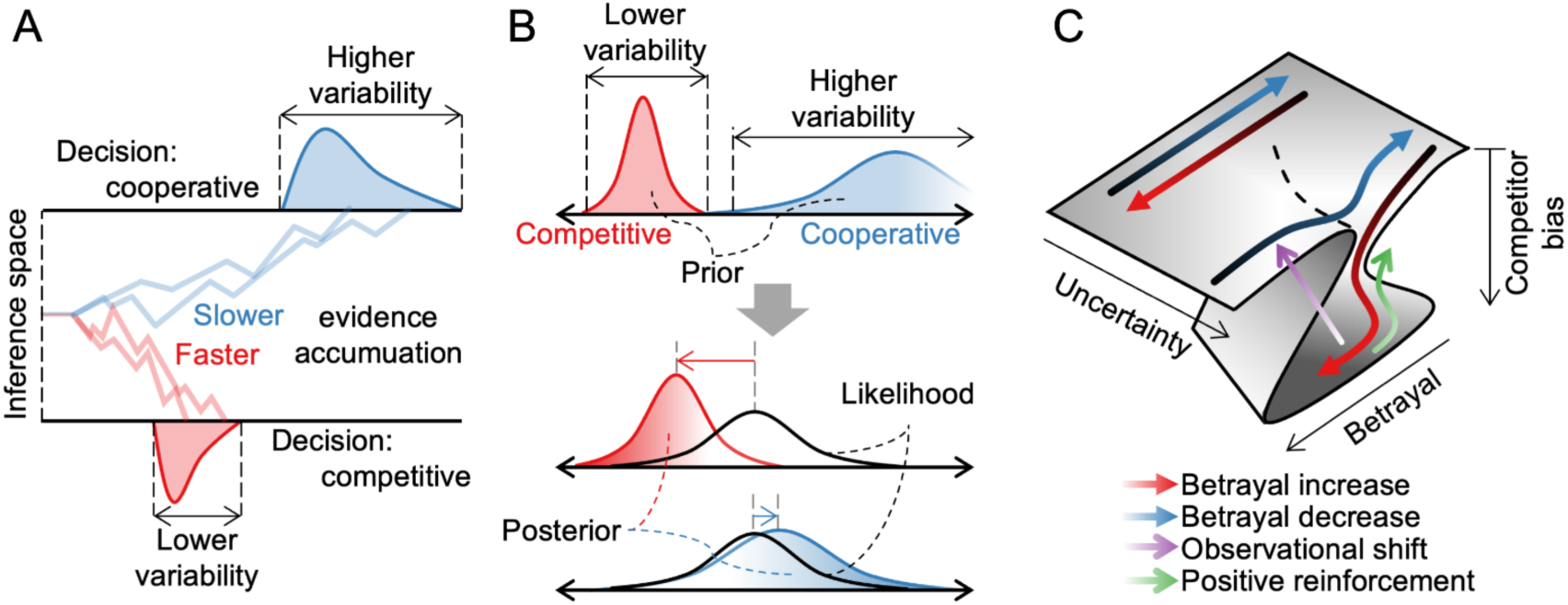
The dynamical systems perspective and alternative viewpoints on hysteresis in social inference. **(A)** Alternative view 1: Drift diffusion model (DDM), where social inference accumulates evidence over time until reaching cooperative or competitive decision thresholds. **(B)** Alternative view 2: Bayesian inference, where history sharpens the strong prior belief that the opponent is a competitor, biasing the posterior distribution. Both models predict less variable decision times or choices. **(C)** Dynamical system perspective: phase diagram showing where social inference stabilizes. In contexts with no uncertainty (extreme Fs), inference shows no bias regardless of betrayal. However, in ambiguous contexts (F≈0.5), the increasing pattern of betrayal biases inference toward perceiving others as competitors, whereas a decreasing pattern does not. Recovery from bias can occur either by gradually reducing betrayal (green arrow) or shifting observation from others to oneself (purple arrow).

Another benefit of the dynamical systems perspective is its ability to better explain the complex temporal dynamics of social cognition. Previous models, such as reinforcement learning based on social prediction error (Apps et al., 2016; Joiner et al., 2017; Koster-Hale & Saxe, 2013; Ruff & Fehr, 2014), often assumed that humans update inferences uniformly from any experience, regardless of the temporal order of previous experiences. However, the same experience can lead to different inferences depending on the order of events—a pattern well captured by hysteresis in dynamical systems. For example, in psychiatry, traditional models struggle to explain why depressive states persist even after stressors have subsided, whereas dynamical systems account for this persistence through hysteresis (Borsboom, 2017; Cramer et al., 2016; Nelson et al., 2017; Scheffer et al., 2024). Hysteresis also illustrates inertia in neurological states that were previously unaccounted for (Friedman et al., 2010; Jirsa et al., 2014). Similarly, we demonstrate hysteresis in social inference; humans continued to suspect betrayal even after it had ceased. Thus, in line with recent advances viewing social interaction through the lens of dynamical systems (Nair et al., 2023; Vinograd et al., 2024), we highlight the need to incorporate the temporal structure of experience into computational models to fully account for how social inference unfolds over time.

Although observation was emphasized as an active mechanism for social inference, this study is limited by the lack of direct manipulation of its influence. Similar to how occlusion in visual attention or first-person perspective constraints influence spatial navigation, direct manipulation of observability could shape social inference. For example, a study demonstrated that manipulating the gaze direction of computer agents altered observers’ inferences about animacy (Gao et al., 2010). More broadly, adjusting observability may help mitigate cognitive biases; individuals with paranoia and alexithymia often exhibit either excessive observation of others or reduced self-observability (Feldmanhall et al., 2013; Finn et al., 2018), and individuals with autism spectrum disorder disproportionately observe non-social information (Dalton et al., 2005; Fortier et al., 2022; Klin et al., 2009). Therefore, future research should investigate whether manipulating observability influences inference and whether inference outcomes, in turn, shape subsequent observational choices. This can be reframed as a fundamental question of whether observational selection can function as a control mechanism for social inference within a closed-loop system. Though challenging, these questions prompt us to move beyond experimental designs that treat social inference as temporally discrete and independent and toward a more naturalistic and closed-loop setting (Maselli et al., 2023; Yoo et al., 2021; Zaki & Ochsner, 2009).

## MATERIALS AND METHODS

### Participants

Data were collected from 54 human participants (31 male and 23 female; age = 22.7±2.3). Forty of them participated in Experiment 1 (23 male and 17 female; age = 22.6±2.2), and the remaining fourteen were in Experiment 2 (8 male and 6 female; age = 23.1±2.8). The Institutional Review Board of Sungkyunkwan University approved the study (IRB 2022-11-009), and all studies followed relevant guidelines. All participants provided written informed consent and were naive to the purpose of the experiment.

### Experimental apparatus

All stimuli were displayed on a 24-inch LCD monitor (S27R350FHK) with a resolution of 1920 × 1080 pixels and a refresh rate of 60 Hz in a dark room. Participants rested their heads on a chin rest at a viewing distance of 60 cm from the monitor. They played the prey-pursuit game (see next section), controlled by a custom MATLAB (MathWorks Inc.) program using the Psychophysics Toolbox (Brainard, 1997; Pelli, 1997). They responded using a Logitech Extreme Pro 3D joystick. The horizontal and vertical eye positions of the right eye were simultaneously recorded using an EyeLink 1000 Plus infrared eye tracker (SR Research) at 1 kHz and later downsampled to 60 Hz for post-hoc analysis. Eye tracker calibration was performed by presenting a circular dot (0.3° in diameter) at nine locations: the four corners, four sides, and the center of the screen. Calibration was repeated if the recorded eye position deviated visibly from the fixation point or if the participant lifted their chin from the chin rest.

### Prey-pursuit game task

Participants played a prey-pursuit game where they had to catch prey as quickly as possible while interacting with an opponent (Figure 2A). Each trial consisted of four phases. In the first phase (inference phase), they watched a video where three characters (player’s avatar, computerized opponent’s avatar, and prey) moved according to their pre-determined algorithms (see the sections below). The prey’s initial position was randomly selected, while the player and opponent avatars were randomly placed within a range from 20° to 20.5°, maintaining an equal distance from the prey. The video ended automatically when either the player or the opponent came within Euclidean distance of 100 pixels of the prey, with an average duration of 2.3 seconds. This was to prevent the opponent or player from revealing hidden intentions too explicitly by catching the prey. The characters were outlined with a black border to prevent confusion with other phases. The video restarted from the beginning if their gaze deviated by more than 20° from the center of the three characters or if they tried to control the joystick. This was to ensure participants paid close attention during the inference phase. Participants were informed that the opponent could be either cooperative, assisting in the hunt, or competitive, intercepting the prey, but were not given details about their exact behavior patterns.

When the item decision phase began, the black outlines of all three characters were removed, and their final positions from the previous inference phase were retained. At the same time, two colored arcs appeared around the player’s avatar (3° apart). Participants were required to move the player’s avatar toward one of the two arcs using a joystick (i.e., a two-alternative forced choice, 2AFC). Reaching the red arc made the opponent’s speed faster in the subsequent phase (boost item), while reaching the gray arc slowed it down (hinder item). The angles at which the two options appeared were fully randomized. The item decision phase lasted until the participant responded, with an average response time of 1.3 seconds.

In the following pursuit phase, all three characters’ initial positions were re-randomized using the same method for the inference phase. From the reset initial position, participants were instructed to control the player’s avatar with a joystick to catch (touch) the prey’s avatar. They had up to 15 seconds to catch the prey, with an average completion time of 7.6 seconds. If they succeeded within 2 seconds, their score in the feedback phase was 100 points and reduced proportionally (e.g., 100 points for catching the prey in 2 seconds, 50 points if caught in 8.5 seconds, and 0 points if not caught in 15 seconds). To prevent participant demotivation, if the opponent caught the prey before the player, the participant received a small portion of the score they would have earned by catching it themselves. Specifically, the original score was multiplied by the opponent’s F value and 0.5. For example, if the player had earned 80 points by catching the prey themselves, they received 8 points when the opponent’s F value was 0.2 (8 = 80 × 0.2 × 0.5). The cumulative score was then proportionally scaled to determine their final experiment monetary compensation.

Participants were given a total of 20 minutes of pursuit time in Experiment 1 and 40 minutes in Experiment 2, excluding the inference phase and item decision periods. This time limit was set to encourage participants to catch the prey as quickly as possible. On average, participants did 216±5 trials in experiment 1 and 505±11 trials in experiment 2.

### Prey algorithm

The prey was designed to achieve two goals: escaping from the opponent and player avatars and staying near the center of the screen to avoid being cornered (Figure 2B). At each time point, the prey selected its next possible 15 locations, evenly spaced 24° apart in polar coordinates and equidistant in Euclidean space from its current position. If any of the 15 possible locations overlapped with a border or another character, that specific location was removed from the options. The change in Euclidean distances from the player and opponent avatars of each possible next position was summed and min-max normalized across all 15 locations (escape cost), and the distance from the screen center was also min-max normalized (centrality cost). The final movement decision was made by weighing the escape and centrality costs at a 6:4 ratio and selecting the position with the lowest combined value. If the prey had visited the same location within the previous two frames, it did not select that position and instead chose the one with the lowest cost among the remaining options. When the opponent was within a distance of 25°, the prey fled at a maximum speed of 1.95°/s (130% of the player’s default speed). If the opponent was farther away, the speed gradually decreased following a sigmoid function, approaching zero beyond 50°.

### Opponent’s avatar algorithm

Similar to the prey’s algorithm, the opponent selected its next position from 15 evenly distributed candidate locations at each time point. The candidate locations were ignored if they overlapped with borders or other characters. The opponent predicted the prey’s subsequent position based on the prey’s algorithm (see above section) for each potential move. Then, the distances to the opponent’s potential position (d_oppo._) and the current player’s position (d_play._) were computed from the predicted subsequent prey position. The experimentally predetermined opponent’s F value, ranging from 0 to 1, determined the opponent’s cost for each potential position (Figure 2C), as follows:

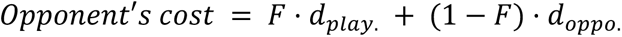

The opponent moves to the position with the lowest cost. This means that when F = 1, the opponent’s movement would position the future prey closer to the player’s avatar, whereas when F = 0, it moves to bring the future prey closer to itself. As a result, from the participant’s perspective, F=1 leads to a shepherding behavior, while F=0 results in a prey-intercepting behavior (see Figure 2E and Supplementary Videos 1 and 2 for examples). These modulations are known to be effective, as previous studies have shown that parametric modulation of physical features can influence perceived intention (Gao et al., 2009). Note that while prior research indirectly modulated intention, we manipulated the computer agent’s latent intention directly.

During the inference phase, the opponent’s speed was 1.65°/s (110% of the player’s default speed). During the pursuit period, if the participant selected the boost item in the item decision period, the speed increased to 2.13°/s (30% increase), whereas if they selected the hinder item, the speed decreased to 1.32°/s (20% decrease).

### Player’s avatar algorithm

To make it difficult to catch the prey without the opponent’s help, the prey’s speed was 1.5°/s, 30% faster than the player’s avatar’s default speed. During the inference period, the player’s avatar moved according to a predefined algorithm, which predicted the prey’s position 750 ms ahead based on its average movement direction over the last five frames and guided the avatar toward that location, with the default speed. This time window of prediction was determined based on previous research suggesting that the brain predicts approximately 750 ms into the future in similar pursuit tasks (Yoo et al., 2020).

During the item decision and pursuit periods, the player’s avatar moved proportionally to the joystick’s force and direction, with a maximum speed when pushed at full strength. Vertical or horizontal joystick inputs that caused overlap with a border or other characters were ignored. To encourage accurate predictions of the opponent’s F value during the item decision period, the player’s speed in the pursuit period was slightly adjusted based on decision accuracy. Specifically, when the opponent’s behavior was predictable (F = 1 or F = 0) and the choice was correct, the player’s speed increased to match the prey’s maximum speed. Conversely, if the choice was incorrect, the speed difference from the prey could increase up to twice the original gap. When the opponent’s behavior was ambiguous (F = 0.5), the player’s speed remained unchanged regardless of the participant’s choice.

### Betrayal and Unexpected-help conditioning

A key manipulation in the experiment was that the opponent’s F value during the inference period differed from the F value during the pursuit period without informing the participants. That is, this change occurred after participants decided whether to boost or hinder the opponent during the item decision period. If the change in F was negative, participants experienced a betrayal condition, whereas a positive change in F led to an unexpected help condition. From the participant’s perspective, the opponent initially appeared to herd the prey toward the player in the betrayal condition but later intercepted it, whereas the opposite appeared to occur in the unexpected help condition. Example trials can be found in Supplementary Videos 1 and 2. In Experiment 1, the betrayal condition was repeated in a block design for 20 trials (with a random variation of ±4 trials) (Figure 2D). The magnitude of F change was randomly set between ±0.35 and 0.45. This value was set so that most participants did not notice the change in *F* while still affecting their pursuit performance. In Experiment 2, the change in F value gradually increased from −0.45 to 0.45 over nine trials (betrayal decreasing phase) or decreased in the opposite direction (betrayal increasing phase) (Figure 6C). These two patterns alternated in a zigzag sequence of increases and decreases. In all cases, F remained within the range of 0 to 1.

### Analysis of item choice and reward

To assess the relationship between participants’ choices and monetary rewards, we binned F values during the inference period into three equal intervals and compared the average reward for each boost and hinder choice (Figure 2F). To assess whether the betrayal and unexpected help conditions affected participants’ monetary rewards, we also calculated the difference in overall rewards between boost and hinder choices (Figure 2G).

To examine whether prior experiences influenced choices, we binned the experimentally given *F* values from the inference period into ten equal intervals and calculated the probability of a boost choice by coding the boost choice as 1. Then, we fitted a cumulative Gaussian curve to capture the psychometric curve of intention inference. In Experiment 1, psychometric curves were computed separately for trials following the betrayal and unexpected help (Figure 3A and Supplementary Figure 1A), and in Experiment 2, the change in F (ΔF) was binned into seven equal intervals (Figure 6D).

To quantify bias in inferences of the opponent’s intention, we coded boost item choices as 1 and subtracted the given F value. Negative values indicated that participants rated the opponent’s intention lower than its actual F value (competitive bias), while positive values indicated a higher rating (cooperative bias). The average bias in intention inferences for male and female participants separately is shown in Supplementary Figure 1B, while the relationship between bias and age is presented in Supplementary Figure 1C. Supplementary Figure 1D presents the trial-averaged inferential bias and its changes as betrayal and unexpected help trials are repeated within the block. Supplementary Figure 1E displays the same data realigned to the moment of block change, as the change occurred randomly around the 20th repetition. In Experiment 2, the change in F (ΔF) was divided into seven intervals, and inferential bias was calculated separately for each interval (Figure 6E).

To measure the internal uncertainty of participants’ inference, the probability of choosing the boost item was converted into Shannon entropy (Figure 3B) as follows;

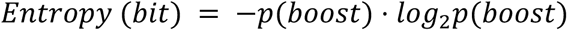

When p(boost) is close to 0.5, entropy approaches its maximum value of 1 bit, indicating maximal uncertainty in choice. When p(boost) is close to 0 or 1, entropy approaches its minimum value of 0 bits, reflecting maximal certainty. Shannon entropy has been used as a measure of internal certainty (Wang et al., 2023), and more broadly, neural encoding of certainty is thought to manifest in choice probability (Kepecs et al., 2008; Meyniel et al., 2015).

### Energy landscape analysis during the inference phase

To derive the structure governing the inference process, we built an autoregressive logistic regression model with two regressors (Figure 3D). The first regressor, *β_t_,* was on a short video clip capturing the trajectories of the three characters. To construct this, we computed the pairwise Euclidean distances among the three characters at each time point over a 1s period, then averaged them within 200-ms bins, resulting in 15 representative feature values (3 characters × 5 time points). The pairwise Euclidean distance was used to reduce confounding in the regression by focusing only on the relative spatial relationships among the prey, opponent, and player, regardless of their absolute positions on the screen. This approach was chosen specifically to avoid using raw coordinates, which could introduce irrelevant variability tied to the layout of the display.

The second regressor, *β_hys,_* was on the scalar change in F (i.e., ΔF, or the amount of betrayal or unexpected help) experienced in the previous trial. Using these two regressors, we predicted whether participants would choose the boost item (coded as 1) or the hinder item (coded as 0) in the following item decision period. The model was trained separately for each participant using all data in experiment 1 from the inference phase, regardless of the betrayal or unexpected help condition. The same method was applied in Experiment 2, except that the betrayal-increasing and betrayal-decreasing phases were trained separately, based on their distinctive patterns shown in Figure 6D-G.

Using the trained weights, we estimated the time series of inferred 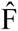 value for each trial. The inferred 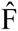 value represents the probability of the participant choosing the boost item, reflecting the strength of their judgment of the opponent as a cooperative helper. We then reconstructed the psychometric curve at each time point using the same method as in Figure 3A, based on the time series of inferred 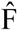 value for the betrayal and unexpected help conditions (Figure 3E). To analyze the meaning of each regressor on intention inference, we conducted simulations by multiplying 0, 1/3, 2/3, or 1 on *β_t_*, or multiplying 0, 1, 2, or 3 on *β_hys_*. We then estimated the psychometric curves based on the inferred 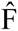 value from the last 1 second before the item decision phase in the same manner as Figure 3E (Supplementary Figure 2). As β_hys_ increases, the difference between unexpected help and betrayal conditions grows, while higher β_t_ steepens the psychometric slope. This suggests that β_hys_ underlies asymmetrical inferential bias while β_t_ governs the inference process during the inference phase.

To reconstruct the energy landscape, we applied the principle that high-energy states exhibit greater temporal variation, while low-energy states show less variation over time. A similar approach is known to successfully reflect the decision space (Wang et al., 2023). First, we divided the trials into three groups based on the given *F* value during the inference phase: low (F< 0.33), medium (0.33 < F < 0.67), and high (F > 0.67). It was further separated into the trials after betrayal and unexpected help in Experiment 1, and into seven equal bins of change in F value (ΔF) in Experiment 2. Second, time points were categorized into twenty equal intervals based on the inferred 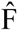 value at each moment, and the average time derivative was computed within each bin (as shown in Figure 3F). Third, the time derivative was integrated across the inferred 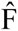 value to estimate the energy landscape of inference. To determine the depth of the basin, we measured the range of energy values, from maximum to minimum (Supplementary Figure 3, left panel). Lastly, we performed min-max normalization on the energy landscape for a fair comparison across participants (Figure 3G, Figure 6H, and Supplementary Figure 6A). Assuming the lowest energy point as the stable fixed point of inference, we fitted first- to third-order polynomial functions to the energy landscape. We then calculated the Akaike information criterion (AIC) values for each fit and estimated the stable fixed point as the minimum of the polynomial function with the lowest AIC value (subfigures in Figure 3G, Figure 6I, and Supplementary Figure 6B). The majority exhibited a U-shape, with a second-order polynomial function providing the best fit. To calculate the landscape’s slope, we split the fitted polynomial function into two parts at the stable fixed point (lowest point). We then applied linear regression to each part to compute the slope. The higher value of the two resulting slopes was defined as the basin’s steepness (Supplementary Figure 3, right panel).

### Gaze comparison analysis

To compare whether participants focused more on the opponent or the player’s avatar, we subtracted the distance from the gaze center to the screen location of the player’s avatar from the distance to the screen location of the opponent at each time point. A negative value indicated that the gaze was closer to the opponent, suggesting a greater focus on the opponent’s movements, while a positive value indicated the opposite. Experiment 1 was analyzed separately for the betrayal and unexpected help conditions (Figure 4B and F, Supplementary Figure 4A), while Experiment 2 was analyzed by dividing the change in *F* (the strength of betrayal or unexpected help) into seven equal intervals (Figure 6F). The gaze comparison value was averaged over the period from 1.5 to 3 seconds after the onset of the pursuit period (Figure 6G). To examine the relationship between observational focus during the pursuit phase and inference in the next trial, we calculated the Spearman correlation coefficient between gaze distance at the opponent (i.e., the strength of negative gaze comparison) and negative biases (perceived F - given F) (Figure 4C). We also calculated the correlation between how long participants gazed at the opponent and their negative bias (Figure 4D). Eye movements exceeding 30° per second were classified as saccades (Supplementary Figure 4B).

### Analysis of control stability

To assess participants’ control stability over their avatar, we calculated jerk strength (Figure 4E). Specifically, we computed the third derivative of the avatar’s trajectory (i.e., jerk), squared it, and then averaged the result. Jerk strength for the prey and opponent’s trajectories was measured similarly, along with the average distance between the gaze location and all three characters. Finally, we computed the partial Spearman’s correlation coefficient between gaze distance to the self’s avatar and jerk strength, controlling for gaze distance to the other characters.

### Task-optimized RNN

To verify whether gaze target shifts causally influence inferential bias, we developed a task-optimized recurrent neural network (RNN) by following the steps (Figure 5A). In the first step, we trained a three-layer Long Short-Term Memory network (LSTM) to predict F value from trajectory sequences. The model consisted of a sequence input layer (input size of 6, which was the x and y coordinates of three characters), an LSTM layer (64 hidden units), and a fully connected output layer (scalar output). It was trained on the inference phase of randomly selected 1000 trials, using the Adam optimizer (learning rate: 0.01, gradient decay: 0.9) with mean squared error (MSE) loss. Gradients were iteratively computed and updated using accumulated values across trials to prevent the influence of any single trial from dominating the learning process. Participants’ responses were not used for training the model.

In the second step, we examined how the neural network adapted to betrayal and unexpected help scenarios. To do this, the model trained in the first step was independently re-trained 1,500 times as follows. The trajectories from the pursuit phase were used as inputs instead of those from the inference phase, as it done in the first step. A total of 200 trials were used, with 100 randomly selected from betrayal trials and the remaining 100 from unexpected help trials, irrespective of participants. The selected set of 200 trials was changed for each re-training cycle within the 1500 training iterations. As a result, the 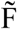 value inferred by the RNN deviated from the given *F* value in the corresponding inference phase. This deviation yielded MSE loss from a negative difference in the betrayal trials (Loss−) or a positive difference in the unexpected trials (Loss^+^). The total loss from 200 trials was computed as follows;

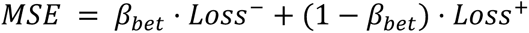

where β_bet_ represents sensitivity to betrayal. When β_bet_=1, the model learns only from means squared error (MSE) generated by negative differences trials (i.e., betrayal), while when β_bet_=0, it learns only from MSE generated by positive differences trials (i.e., unexpected help). β_bet_ was initially set as neutral (=0.5). Next, with the recurrent and output layer weights fixed, the input layer weight and β_bet_ were optimized using the same procedure as in Step 1, based on total MSE. This was to examine whether the RNN mimicked the observational focus shift observed in humans in the context of betrayal or unexpected help. For each of the re-learning iterations, the optimization process was repeated until β_bet_ reached an extreme value (either above 0.9 or below 0.1). The changes in β_bet_ across epochs are shown in Figure 5B, and the given F values from the inference phase used for training are shown in Figure 5C.

To assess potential overfitting in training, half of the trials were used as the training set (50 trials for the betrayal type and another 50 trials for the unexpected help type). The remaining trials, set aside separately, were used as the test dataset. The accuracy (1 - total loss) of both the training and test sets across epochs revealed that the test dataset’s accuracy slightly declined from the 65th epoch, but the drop was negligible (∼0.5%) (Supplementary Figure 5). Additionally, the tendency to categorize into two networks was already evident from the 65th epoch. By the 65th epoch, nearly all re-learning iterations had already reached the extreme value of β_bet_. Therefore, the current results were not considered to be due to overfitting.

We examined the step-2 RNN in several ways. Firstly, the inferential bias of the RNN model was measured (Figure 5D). To achieve this, inference phase trajectories from step 1 training were provided as input again without additional weight optimization (i.e., Forward pass test). The bias was calculated by subtracting the actual F value from the inferred 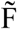 value, following the same approach used in the choice analysis (shown in the subfigure of Figure 3A). We further tested whether the changes in the activation of each unit were correlated with the inferential biases of the model (Figure 5E, left panels). For interpretability, the sign of the weight change was reversed if it showed a negative Spearman correlation with the RNN’s inferential bias. This was to ensure that the activation increase of an RNN unit consistently signaled a cooperative bias. Only units with a p-value lower than 0.05, after FDR correction, were considered significant (Figure 5E, right). We also compared the weight changes associated with the opponent’s position and the player’s position (Figure 5F). A positive change indicated a shift toward a cooperative bias, while a negative change indicated a shift toward a competitive bias.

In the third step, we examine the effect of observational focus shifts on sensitivity to betrayal (Figure 5G). To test this, we reversed the sign of the weights corresponding to the opponent’s and player’s avatar positions in the input layer. All other learnable parameters remained unchanged, except for β_bet_, which was re-optimized from its pre-reversal value. Since the number of parameters to be learned has decreased, the learning rate was adjusted to 0.05. The changes in β_bet_ across epochs are shown in Figure 5H, and the portion of β_bet_ shifting from one extreme to the other (i.e., 0.1 → 0.9 or 0.9 → 0.1) is shown in Figure 5I.

### Statistical test

All pairwise comparisons between two values were conducted using two-tailed, independent t-tests. To examine interactions among three variables, we conducted a repeated measures ANOVA. We conducted a linear trend analysis to evaluate how responses changed consistently across trials. For time series data, to account for the multiple comparison problems, we applied a nonparametric, cluster-based permutation approach to correct the p-value criterion for the two-tailed, one-sample t-test using MATLAB’s statistical toolbox (Maris & Oostenveld, 2007).

---

**Supplementary Figure 1.**
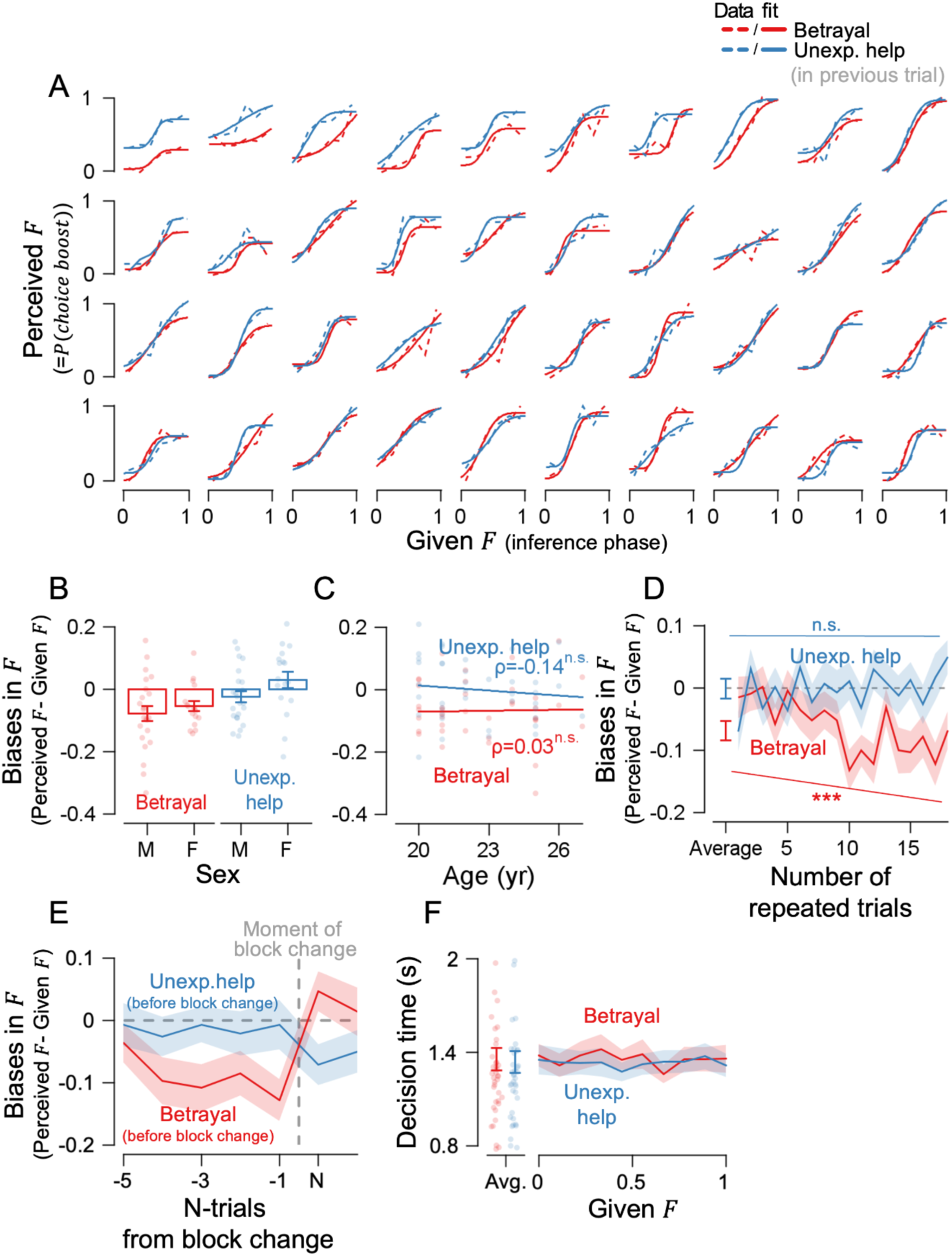
Rejection of possible confounds in item decision of Experiment 1. **(A)** The psychometric curves of F values for the forty participants. The solid line represents the cumulative Gaussian fit, while the dashed lines indicate data. **(B)** Biases in the inference are grouped by sex. **(C)** No correlation was found between participants’ age and biases in the inference. **(D)** Negative biases in perceived F value increased with repeated betrayal but did not after unexpected help. Stars indicate statistical significance (***, p < 0.001; n.s., not significant). **(E)** Biases in perceived F value re-aligned at the moment of block change. Colors indicate the block type before the block change occurred. **(F)** No difference in decision times during the item decision phase between betrayal and unexpected help conditions. Error bars and shaded ribbons represent ±1 SEM, and dots indicate individual participants.

**Supplementary Figure 2.**
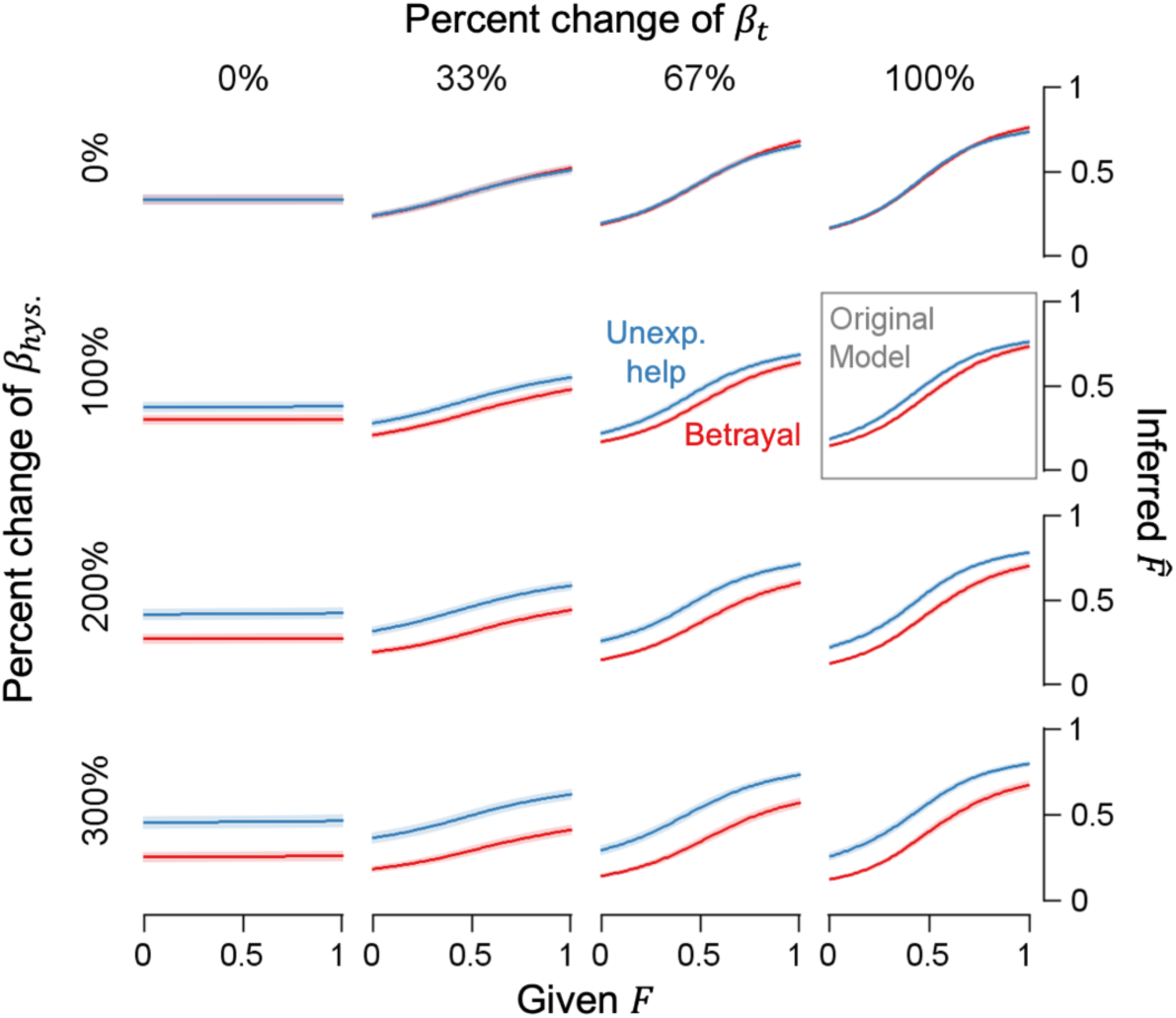
Simulation of parameters in the auto-regressive logistic regression model. The weight on the experience of betrayal or unexpected help (β_hys_) and the weight on the three avatars’ trajectories (β_t_) were parametrically amplified or attenuated. Psychometric curves based on logistic regression model data capture inference during the last second before the item decision phase. Changes in β_hys_ influenced the overall baseline difference between the betrayal and unexpected help conditions, while changes in β_t_ affected the inferential bias in a given context. Shaded error bars represent ±1 SEM.

**Supplementary Figure 3.**
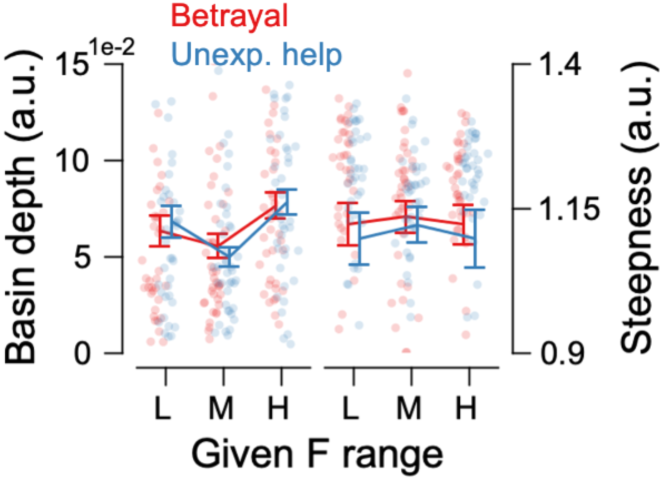
Certainty of inference did not differ across conditions in the energy landscape. The depth (difference between minimum and maximum values) and steepness (slope) of the inference energy landscape were similar across the basins of betrayal and unexpected help. Error bars represent ±1 SEM.

**Supplementary Figure 4.**
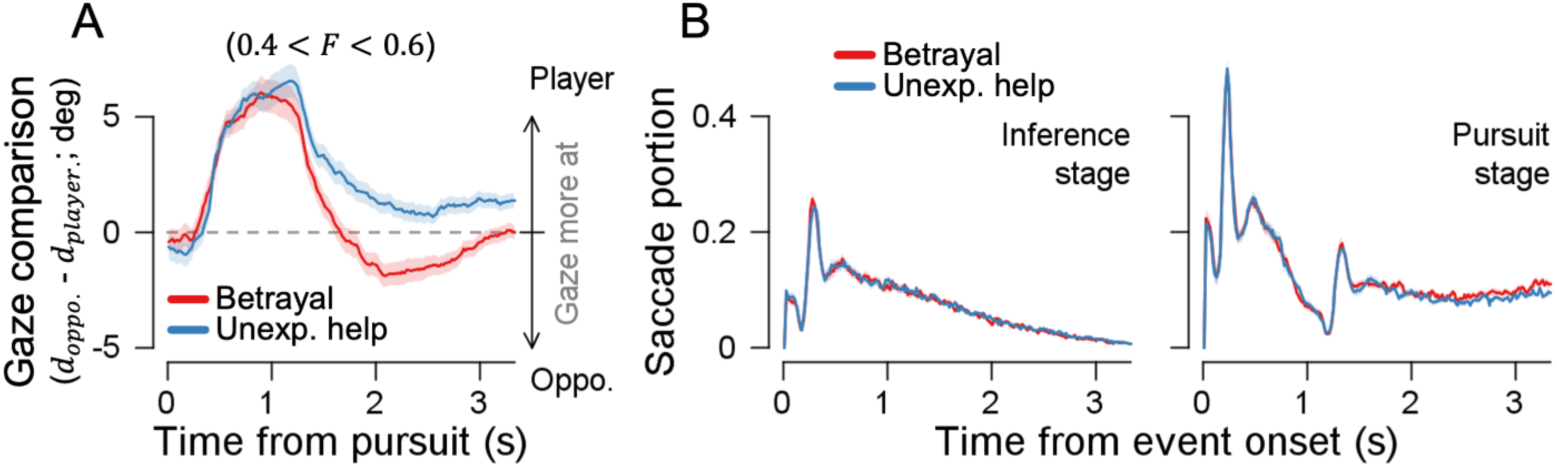
Rejection of possible confounds in observation results of Experiment 1. **(A)** Comparison of player and opponent gaze during the pursuit period, restricted to trials with matched F values (0.4 < F < 0.6). **(B)** The proportion of saccades (eye movements with speed over 30°/s) during the inference phase and pursuit. Shaded error bars represent ±1 SEM.

**Supplementary Figure 5.**
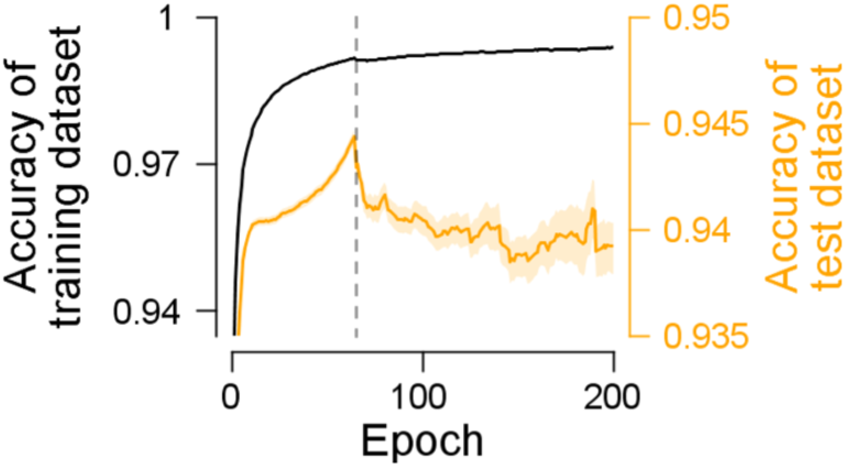
Accuracy of F value prediction by the task-optimized RNN during step 2. Black: training dataset (half of the dataset). Yellow: the test dataset (the other half). Shaded error bars represent ±1 SEM.

**Supplementary Figure 6.**
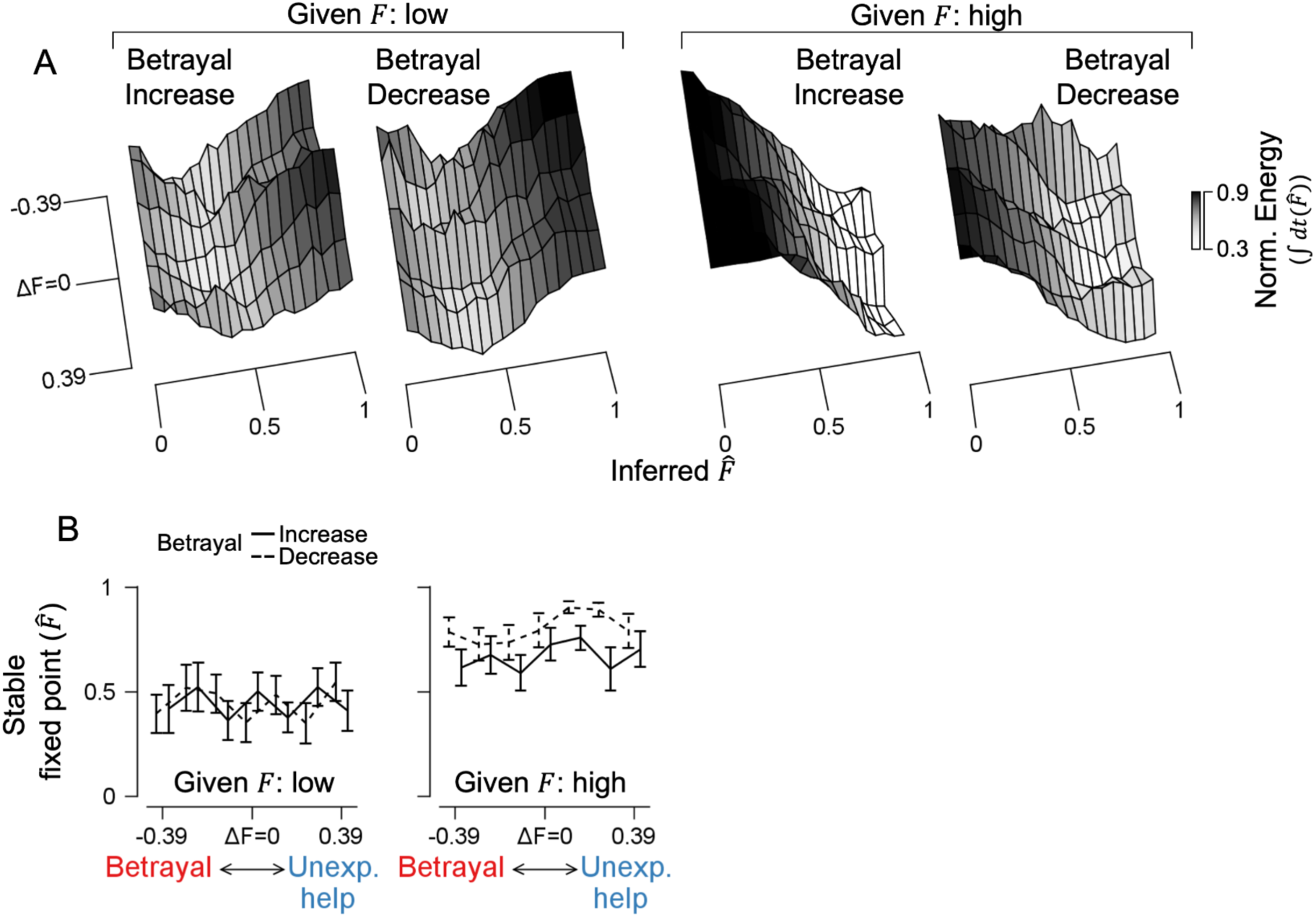
Reconstructed energy landscapes of inference. Separate energy landscapes are shown for trials with low (F < 0.33) and high F value (F > 0.67) (A), along with their corresponding stable fixed points (lowest points of the basins) in the phase diagram (B). Error bars represent ±1 SEM. Solid lines indicate the increasing betrayal phase, while dashed lines represent the decreasing phase.

**Supplementary Video 1. Example of betrayal scenario in Experiment 1.** After the inference phase, the participant selected the boost item but subsequently lost the prey to the opponent during the pursuit phase. For illustrative purposes, the opponent’s F value was altered from 1 during the inference phase to 0 in the pursuit phase to depict the most extreme behavioral shift. In the actual experiment, however, the maximum change in F (ΔF) was −0.45.

**Supplementary Video 2. Example of unexpected help scenario in Experiment 1.** During the pursuit phase, the participant captured the prey with ease as the opponent drove it toward the player. For illustrative purposes, the opponent’s F value was altered from 0 during the inference phase to 1 in the pursuit phase to depict the most extreme behavioral shift. In the actual experiment, however, the maximum change in F (ΔF) was +0.45.

## Acknowledgment

This research was supported by RS-2023-00211018 (SS and SBMY). The authors declare no competing financial interests.

## Author contributions

Conceptualization, SS and SBMY; Data Collection, SS; Formal Analysis, SS; Writing—Original Draft, SS and SBMY; Writing—Review and Editing, SS and SBMY; Funding Acquisition, SBMY

## Data and code availability

The codes and datasets generated and/or analyzed during the current study are available at https://github.com/SangkyuSon/socialObservationHysteresis. Correspondence and requests for materials should be addressed to SS and SBMY. The data could be shared upon a reasonable request to the corresponding authors.

